# Structural pathway for class III PI 3-kinase activation by the myristoylated GTP-binding pseudokinase VPS15

**DOI:** 10.1101/2023.09.28.559894

**Authors:** Annan S. I. Cook, Minghao Chen, Xuefeng Ren, Shanlin Rao, Samantha N. Garcia, Ainara Claveras Cabezudo, Anthony T. Iavarone, Gerhard Hummer, James H. Hurley

## Abstract

The class III phosphatidylinositol (PI) 3-kinase complexes I and II (PI3KC3-C1 and -C2) are central to the initiation of macroautophagy and endosomal maturation, respectively. Through three-dimensional classification of a large cryo-EM dataset of human PI3KC3-C1 bound to the small GTPase RAB1A, we were able to map the structural pathway of enzyme activation. The inactive conformation is stabilized by an *N*-myristoyl modification of the pseudokinase (PK) subunit VPS15. The *N*-myristate is sequestered in the N-lobe of the VPS15 PK domain, which stabilizes a series of interactions whereby VPS15 sequesters and blocks the catalytic and membrane binding units of the VPS34 lipid kinase. In the activated conformation, the *N*-myristate and the VPS34 lipid kinase domain are liberated to interact with membranes and catalyze PI3P formation. The VPS15 PK domain contains a unique Arg at the gatekeeper position and binds tightly to GTP. GTP binding structurally stabilizes the *N*-myristate “in” conformation, which promotes the inactive conformation. This pathway provides a general mechanism for PI3KC3 activation in autophagy and endosome biogenesis and a roadmap for their pharmacological upregulation.

---

Macroautophagy (hereafter autophagy) is the primary eukaryotic catabolic response to amino acid starvation, and the main cellular mechanism for clearing toxic aggregates and dysfunctional organelles^1^. Deficits in autophagy are implicated as drivers of many human diseases^2^, as most clearly illustrated by the Parkinson’s disease genes Parkin and PINK1, which initiate damage-induced mitophagy^3^. Autophagy is initiated by the concerted action of several core complexes^4^, including the phosphatidylinositol 3-kinase class III complex I (PI3KC3-C1). PI3KC3-C1 is a heterotetrameric complex comprised of one subunit each of the VPS34 lipid kinase, the regulatory scaffold VPS15, BECN1, and the autophagy specific subunit ATG14^5–8^. The related complex II (PI3KC3-C2) which functions in both endosome maturation and autophagy, shares the three subunits VPS34, VPS15, and BECN1, and has the same overall architecture. In PI3KC3-C2, UVRAG replaces ATG14 as the fourth subunit^5^. PI3KC3-C1 produces phosphatidylinositol (PI) 3-phosphate (PI3P) on the phagophore, which is the precursor to the autophagosome. PI3P recruits WIPI proteins, which are in turn critical for all subsequent steps in autophagosome expansion^9^. In endosome maturation, PI3P generated by PI3KC3-C2^10,11^ recruits the early endosome fusion^12^ and ESCRT machinery^13^ and other essential factors. In both the autophagic and endosomal pathways, PI3KC3 activation is a decisive, committing a membrane to its maturation.

Because of their central role in autophagy and endosomal sorting, the structure and dynamics of PI3KC3-C1 and -C2 complexes have been intensively studied^14–21^. Yet a near-atomic picture of the activating transition has been elusive, frustrated by the dynamic character of the complex. VPS34^KD^ is highly mobile relative to the rest of PI3KC3-C1^14,16^, and its mobility is necessary for function^16^. This has made it challenging to determine the structures of conformational substates at high resolution. Recent advances in the resolution of the apo-PI3KC3-C1 complex^21^ and the insight that the small GTPase RAB1A binds and recruits PI3KC3-C1^20^ have now allowed us to attain a local resolution of nearly 2.3 Å in the best instance, and to classify and refine inactive, transitional, and active conformations, defining the major conformational states along the PI3KC3 activation pathway. We uncovered a large conformational change in the VPS34 kinase domain (VPS34^KD^) relative to the VPS15 pseudo-kinase domain (VPS15^PKD^). We discovered that an Arg residue at the gatekeeper position of VPS15^PKD^, unique in the kinome, confers GTP binding. We were able to map the position of the covalent *N*-myristoyl modification of VPS15. This high-resolution structural insight affords us a picture of the activation cycle of VPS34 as regulated by a previously known role for *N*-myristoylation and the previously unknown binding of GTP to VPS15^PKD^.

## Cryo-EM structure of PI3KC3-C1 in complex with RAB1A

We reconstituted (Extended Data Fig. 1) and determined the structure of the GTP-locked RAB1A(Q70L)-PI3KC3-C1 complex by cryoelectron microscopy (cryo-EM) (Fig. 1a,b, Extended Data Fig. 2). Extensive 3D classification of a 19,000 movie dataset (Extended Data Fig. 2) allowed us to determine the structures of three 3D classes that appeared to represent distinct states in the activation pathway. In the most populated 3D class, determined at nominal resolution of 2.7 Å, VPS34^KD^ is not visualized. We refer to it as the “transitional” structure since it is neither evidently active nor inactive. The overall architecture resembles that of yeast apo-PI3KC3-C2^15^ and apo-PI3KC3-C1^21^ (C_α_ r.m.s.d. = 2.7 Å). RAB1A binding induced the VPS34 helical hairpin (VPS34^HH^) to rotate inward and upward, moving by up to 5.5 Å relative to apo-PI3KC3-C1 to accommodate the RAB1A switch-II motif (Extended Data Fig. 2). This leads to a contraction of the complex along its long axis by 7 Å, a rotation of the VPS15 WD40 and helical linker domain by 5 degrees, and a global bending of the complex resembling a longbow pulled under tension (Extended Data Fig. 3).

**Fig. 1.**
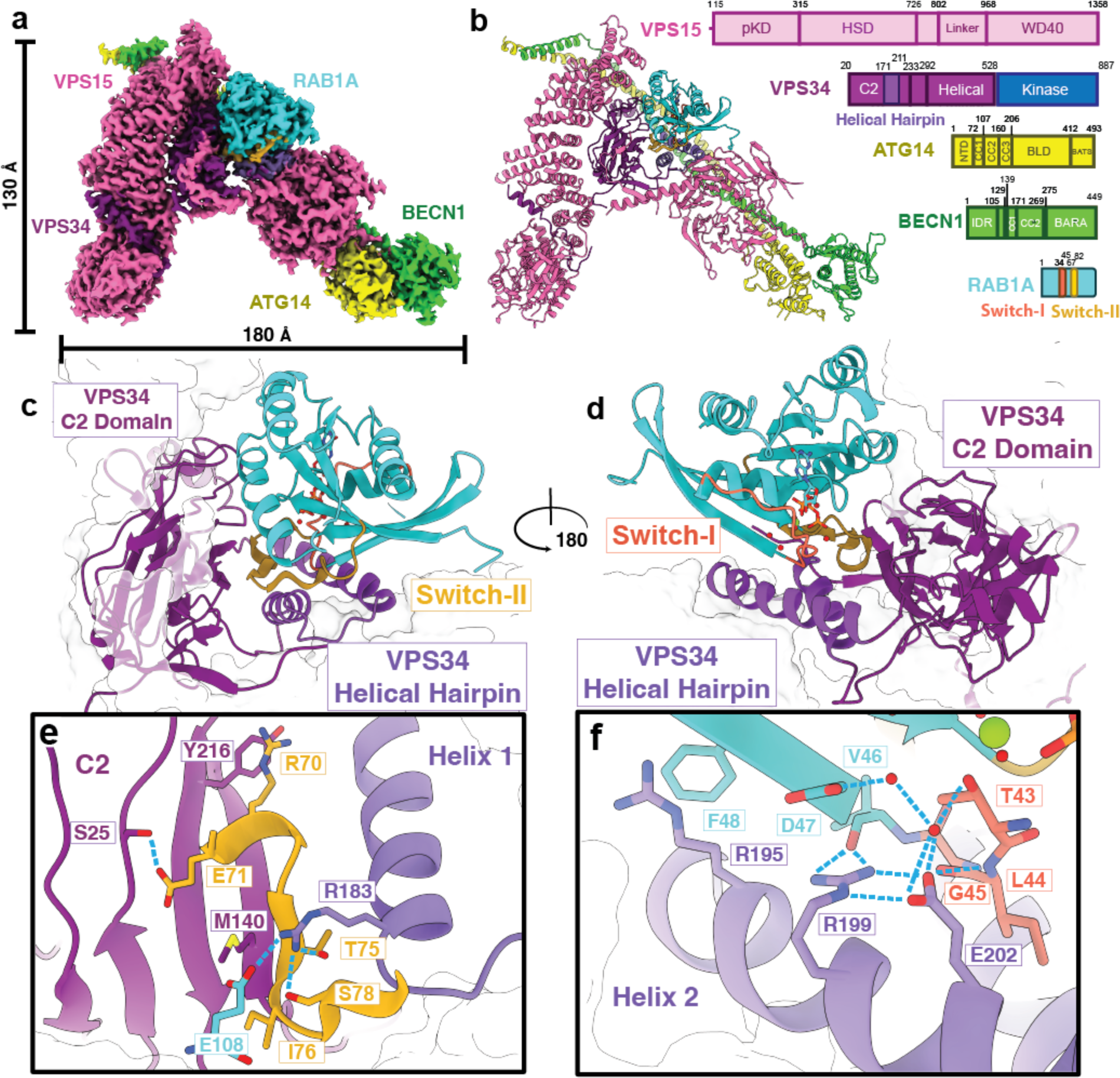
Structure of the RAB1A-PI3KC3-C1 complex. a. Cryo-EM reconstruction of the RAB1A-PI3KC3-C1 and the associated atomic model. b. Domain organization of the complex. Colors correspond to (a). c. Close up of the RAB1A interface with VPS34 C2 domain. The RAB1A switch-II motif (gold) and the HH (light purple) are indicated. d. 180 degree rotation of the RAB1A C2 domain interface. The RAB1A switch-I motif is indicated (orange). e. Inset, close-up of the switch-II motif contacts with the C2-HH cleft. The switch-II helix adopts an extended conformation, forming multiple hydrophobic and polar contacts with HH helix 1 and the C2 β-sheet core. f. Close up of the switch I interface with HH helix 2. The Arg199 and Glu202 form a hydrogen bond network with the switch-I and interswitch domains.

RAB1A binds to PI3KC3-C1 *via* the cleft between the VPS34 C2 domain (VPS34^C2^) and VPS34^HH^ (Fig. 1c,d) *via* the GTP-dependent switch-II region of RAB1A (Fig. 1c). RAB1A switch-I and the interswitch β-strand contacts the first α-helix of VPS34^HH^. The local resolution of the density reached 2.5 Å in this region, and we were therefore able to visualize ordered water molecules near the GTPase reaction center (Extended Data Fig. 4). VPS34^HH^ Arg199 and Glu202, two residues from the REIE motif essential for the RAB1A-PI3KC3-C1 interaction^20^, form a salt bridge and hydrogen bond cage with Asp47 of the RAB1A interswitch motif. VPS34 Glu202 forms a salt bridge with Arg199, which in turn hydrogen bonds to the backbone carbonyl of RAB1A at Asp47. Asp47 of the RAB1A interswitch motif coordinates a pair of water molecules together with VPS34^HH^ Glu202. Glu202 also hydrogen bonds with the amide of the switch-I backbone at RAB1A Gly45, while the water coordinated between VPS34 Glu202 and RAB1A Asp47 contacts switch-I Thr43. Thr43 participates in the octahedral coordination of the Mg^2+^ in the GTP pocket and becomes disordered in the GDP state. The N-terminal and GTP-proximal portion of switch-II is in essentially the same conformation as seen in other RAB1A:effector complexes^22^, while the C-terminal portion from Thr75 to Ser78 diverges to accommodate the first VPS34^HH^ helix. This interaction network explains specificity for the RAB1A GTP-bound state.

## Autoinhibition is driven by the ordered *N*-myristate of VPS15

Refinement of a second 3D class allowed us to build a model of the VPS34^KD^ interface with the VPS15 pseudo-kinase domain (VPS15^PKD^) with residue-level detail (Fig. 2a, b). This conformation resembles those previously observed at lower resolution in complex with NRBF2^17^ and as bound to liposomes^20^. The lower resolution of the previous structures precluded mechanistic interpretation in side-chain level detail. In this conformation and at the newly attained resolution, we observed that the VPS34^KD^ catalytic loop containing the critical DFG motif is sequestered by the VPS15^PKD^ N-terminal region, and the VPS34^KD^ activation loop (A-loop) is in turn sequestered by the VPS15^PKD^ phosphate-binding loop (P-loop) (Fig. 2c). The inaccessibility of the DFG motif and A-loop of VPS34 in this conformation is incompatible with kinase activity. Therefore, we denote this conformation as “inactive”. We observed a density feature in a pocket formed from bulky hydrophobic residues of VPS15^PKD^ P-loop Phe37 and Phe38 and A-Loop Tyr185 and Phe186, Phe55, and VPS15^PKD^ αC Leu64, Tyr67, and Leu71. This density is contiguous with the polypeptide density for the N-terminus of VPS15 (Fig. 2d). VPS15 is known to be *N*-myristoylated^23^, and the density could be entirely accounted for by modeling it as an *N*-myristoyl modification of Gly2 (Fig. 2d). To confirm that this post-translational modification was present in our sample, we performed trypsin digestion coupled to LC-MS and identified a peptide corresponding to the myristoylated N-terminal 29 residues of VPS15 (Extended Data Fig. 5). The VPS34 A-loop Arg798 and VPS15 P-loop make hydrogen bonds to the main-chain of the *N*-myristoyl Gly2 itself, and adjacent residues, including Ile8 (Fig. 1e). These observations establish the structure of the inactive conformation of PI3KC3-C1, the ordered binding of the VPS15 *N*-myristate, and show how the ordered binding of the *N*-myristate in the VPS15^PKD^ helps lock VPS34^KD^ in an inactive conformation.

**Figure 2.**
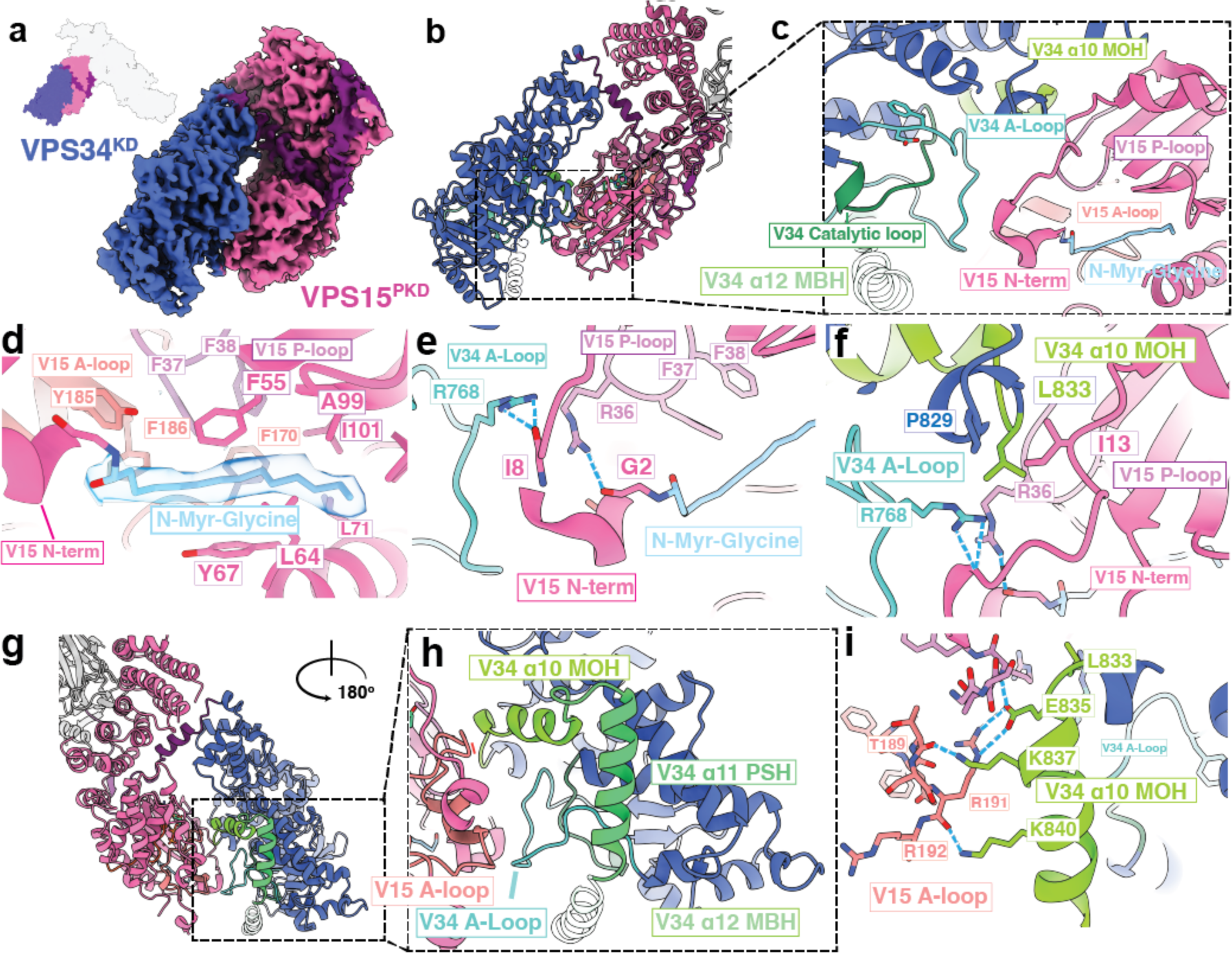
Structure of the VPS34^KD^-VPS15^PKD^ interface and ordered myristate in the inactive conformation. a. Focused view of the Cryo-EM reconstruction of the VPS34 kinase domain in the inactive state. The inset at upper left shows the context of this VPS34^KD^-VPS15^PKD^ interface in the full structure. b. Model of the kinase domain derived from the map. c. Close-up of the interface between the VPS34 A-loop and the VPS15 N-terminal domain. d. The *N*-Myristoyl-Glycine is sequestered in a hydrophobic pocket lined with bulky hydrophobic residues from the VPS15 activation loop, the VPS15 P-loop, and the C α-helix domain. Density is shown for the *N*-myr-Gly at a contour of σ = 7.0 e. The interaction between the VPS34 A-loop and the VPS15 N-Terminal Domain. Hydrogen bonds are denoted with black dashed lines. f. Hydrophobic contacts with VPS15 NTD lock VPS34 MOH in a position that blocks membrane interaction by VPS34^KD^. g. Rotated view of the VPS34 inactive interface. h. Inset view of the sequestration of VPS34 MOH and PSH by VPS15^PKD^. The VPS34 MBH is indicated in transparent ribbon form as its position was inferred from the orientation of the other secondary structural features of the kinase domain and not visualized in the cryo-EM density. i. VPS34 MOH forms several hydrogen bonds (black dashes) with the carbonyl oxygens of VPS15 A-loop and P-loop, as well as between VPS34 Glu835 and Arg191.

The C-terminal helix of VPS34, Kα12, embeds into the membrane and is essential for catalysis^24^. Because of its key role in function and membrane-binding geometry, we refer here to Kα12 as the VPS34 membrane binding helix (MBH). MBH is positioned in part by the immediately preceding helices Kα10 and Kα11 (Fig. 2g,h), and because of its role in positioning the MBH, we refer to Kα10 as the “MBH orientation helix” (MOH). The VPS15^PKD^ A-loop interacts extensively with VPS34 MOH (Fig. 2i), reaching across the narrow gap between VPS34 and VPS15. VPS34 MOH Glu835 interacts with VPS15 A-loop Arg191 and the VPS15 P-loop main chain amides of Thr35 and Arg36. VPS34 MOH residues Lys837 and Lys840 donate hydrogen bonds to the carbonyl oxygens of VPS15 A-loop Thr189, Arg191, and Arg192. VPS15 N-terminal region Ile13 forms hydrophobic contacts with VPS34 MOH Leu833 and the preceding Pro829. In essence, the VPS15 N-terminus and A- and P-loops form an extensive web of contacts that sequesters the MOH, resulting in the misorientation of the MBH and making it impossible for VPS34^KD^ to engage with its lipidic substrate. Thus, the VPS15 *N*-myristate “in” state blocks both the catalytic and membrane binding elements of VPS34^KD^.

## Liberation of the VPS34^KD^ catalytic and membrane-engagement motifs in the active conformation

Further 3D classification of the same dataset captured a third 3D class, representing a distinct and previously unresolved conformation of VPS34^KD^. We refined this state to a nominal overall resolution of 3.3 Å, although density for VPS34^KD^ and VPS15^PKD^ was of lower quality, which we attributed to a strong preferred orientation of this state, and to increased mobility in the VPS34^KD^ (Extended Data Fig. 2). The local resolution for VPS34^KD^ of 5 Å sufficed to place VPS34^KD^ as a rigid body. Relative to its position in the inactive state, VPS34^KD^ rotated by 140 degrees (Fig. 3a-c, Supplementary Movie 1), breaking all of the VPS15^PKD^ contacts previously visualized in the inactive state. A cluster of Arg residues, including Arg36 of the P loop, and Arg191 and Arg193 of the VPS15 A-loop, formed completely new interactions with VPS34 Kα11 (Fig. 3d, e). Because this helix occupies the cleft in VPS15^PKD^ that would be the substrate binding pocket in a conventional kinase, we term helix Kα11 the “pseudo substrate helix” (PSH). To assess the role of the PSH-VPS15^PKD^ interface in favorably positioning VPS34^KD^ for activity, we made two VPS15 mutant constructs, R191D/R193D and R36D. Despite the possibility of offsetting effects on the inactive conformation, reconstituted complexes containing these mutations were significantly reduced in their lipid kinase activity (Fig. 3f), consistent with their stabilizing roles in the structure of the active conformation.

**Figure 3.**
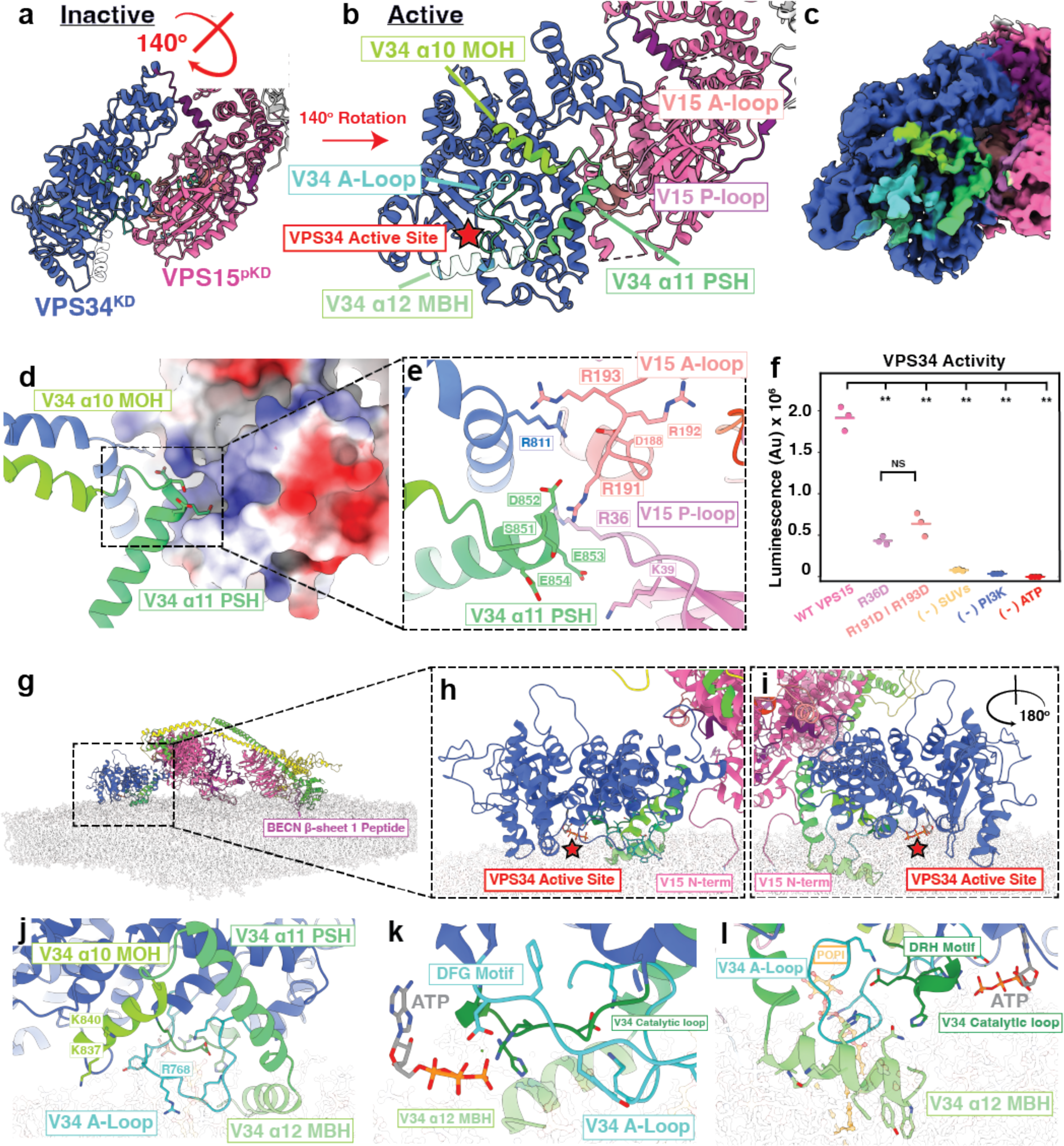
Structure of the VPS34 active state and MD simulation of its membrane-bound geometry. a. Inactive state structure shown for reference. The VPS34^KD^ rotates by 140° upon activation, as depicted by the arrow pointing to b. b. The model of the kinase domain in the active state. Key secondary structural features are indicated with labels in the color corresponding to the secondary structural element. The active site of the kinase is indicated with a red star. c. Density obtained from focused refinement of the VPS34^KD^-VPS15^PKD^ region in the active conformation. d, e. Overview and close-up view of novel VPS34^KD^-VPS15^PKD^ contacts specific to the active conformation. f. Lipid kinase activity of VPS15 mutations in the active site contacts as measured by luminescence assay. The results of three biological replicates were compared by one-way ANOVA to test for significance, and then a post-hoc T-test with the Bonferonni parameter applied was used for pairwise comparisons. g. MD snapshot of the active PI3KC3 complex I engaged with a membrane. h, i. Detailed views in two orientations of A-loop and catalytic loop conformations from snapshot shown in (g). j. Close up view of the VPS34 MOH and A-loop residues that switch interactions between the inactive to active states. k, l. ATP from the MD simulation bound to the VPS34^KD^ in the orientation of (g-i). The catalytic loop and ATP molecule sit at the appropriate height to recognize and orient the substrate PI to facilitate phosphotransfer.

To address the implications of this newly identified conformation for membrane binding, we performed all-atom molecular dynamics (MD) simulations of PI3KC3-C1 on a phospholipid membrane with a phagophore-like membrane composition (Fig. 3d-f). The complex was docked onto the bilayer with VPS34 MBH in direct contact with the membrane, and the resulting geometry was stable during the simulation. As compared to the cryo-ET structure of an inactive PI3KC3-C2 bound to liposomes^20^ through the aromatic tip of the BECN1 BARA domain alone, membrane contact is far more extensive. A BECN1 β sheet-1 peptide near the N-terminus of the BARA domain is known to unfold and participate in membrane binding^18^, yet in previous models of the docked complex, it was too far from the membrane to insert. In the MD simulation of the active complex, this part of BECN1 is in immediate contact with the membrane (Extended Data Fig. 6a). Thus, the newly identified active conformation not only places the VPS34 active site in productive contact with the membrane, but also rationalizes other existing data on PI3KC3-C1 membrane docking that was incompatible with previous models for the membrane interaction.

## VPS15 is a GTP-binding pseudokinase

The 3.3. Å structure of apo-PI3KC3-C1^21^ contained density for a nucleotide molecule in the VPS15^PKD^ corresponding to the ATP binding site of active protein kinases, but that structure lacked the resolution required to unambiguously identify the nature of the nucleotide. Local refinement of the VPS15^PKD^ -VPS34^KD^ of the RAB1A bound complex yielded a local resolution range of 2.3-2.6 Å for the nucleotide pocket (Extended Data Fig. 4). The quality of the density was sufficient to place every side chain that contributed to nucleotide recognition, including associated ordered water molecules. The gatekeeper position responsible for nucleotide specificity in kinases^25^ is Arg103 in VPS15. In functional kinases, which utilize ATP, the gatekeeper is almost invariably hydrophobic^25^. VPS15 homologs are unique among kinases and pseudokinases in containing an Arg at the gatekeeper position (Fig. 4d; Extended Data Fig. 7). Our map revealed both the position of Arg103, and the overall shape and hydrogen bonding network of the nucleotide (Fig. 4a, b). The shape of the purine base density and the position of the Arg103 guanidino group are incompatible with placement of ATP in the density. However, GTP fit the density perfectly, and N5 and O6 of the guanine base are ideally positioned to accept hydrogen bonds from the Arg103 guanidino moiety (Fig. 4b). To confirm the identity of the nucleotide, we denatured PI3KC3-C1 purified from HEK 293 cells and eluted the bound molecule via ion-pair reversed-phase acetonitrile gradient HPLC. In confirmation of the EM density map and hydrogen bonding geometry prediction, the bound nucleotide eluted at the same position as the GTP standard (Fig. 4c), whilst ruling out ATP, and ADP. We further characterized the nucleotide that was released by electrospray ionization MS (Extended Data Fig. 5). This directly confirmed that the dominant nucleotide present was GTP, with no detectable ATP or ADP. The MS analysis did reveal the presence of a lesser proportion of GDP, consistent with the likelihood that VPS15^PKD^ is capable of slowly hydrolyzing GTP.

**Figure 4.**
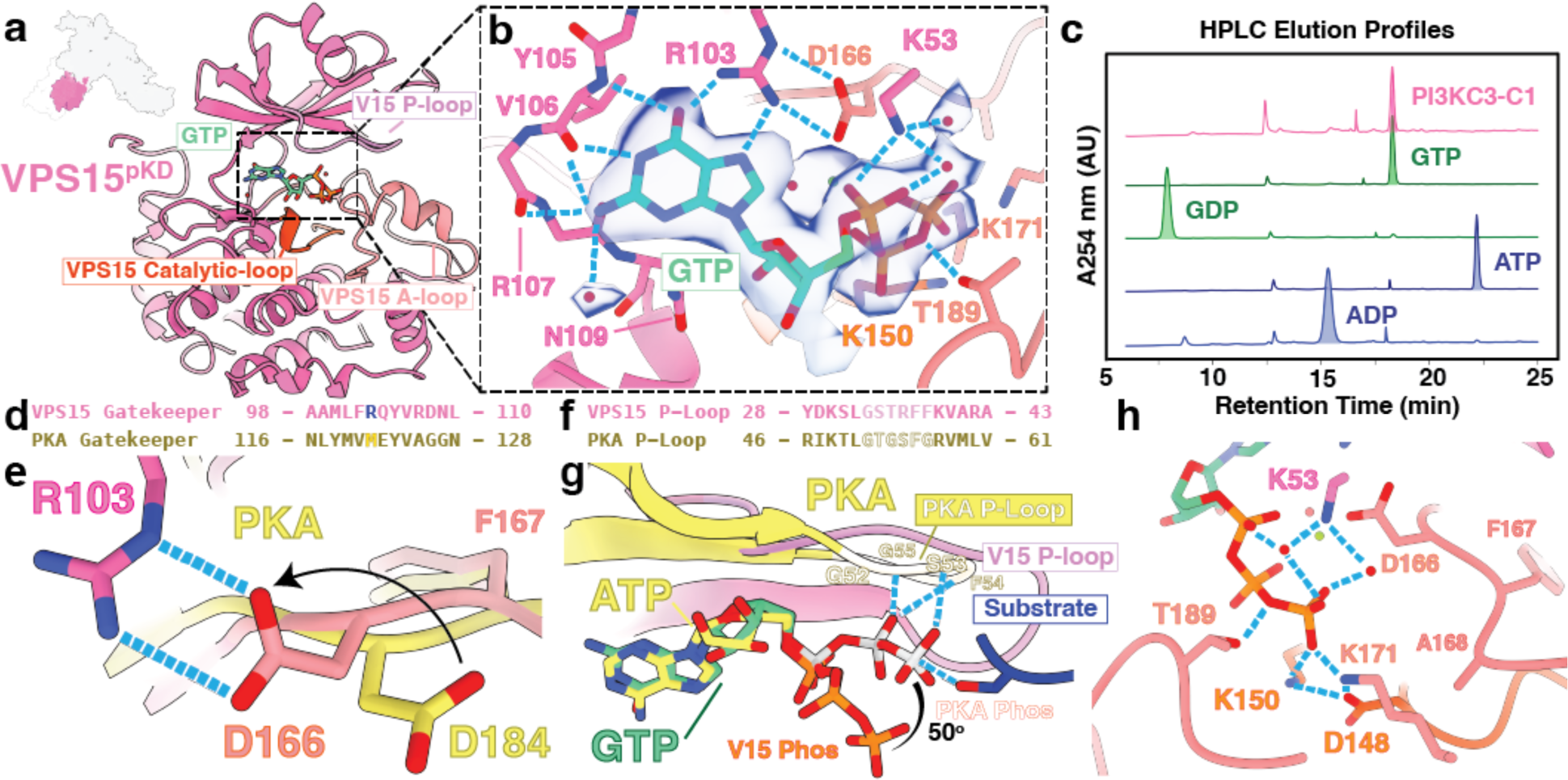
VPS15 is a GTP binding pseudokinase. a. The overall architecture of the VPS15^PKD^. Broadly conserved secondary structural motifs are indicated with labels in the color of each motif. GTP is indicated in green. b. Cryo-EM density (σ = 7) of the bound nucleotide is shown. The hydrogen bonds with Arg103 donate hydrogen bonds to the N5 and O6 groups on the guanine ring. c. HPLC elution profile results. Material eluted from a denatured PI3KC3-C1 sample (pink) is shown to be retained at the same time as the GTP standard. Other nucleotides are shown in the indicated colors. d. Local sequence alignment with *<u>Mus musculus</u>* protein kinase A (PKA) showing the substitution of Arg at the gatekeeper position, indicated in blue. e. The gatekeeper Arginine donates a pair of hydrogen bonds to the DFG Asp, rotating the sidechain away from the normal catalytic orientation shown in the *<u>M. musculus</u>* PKA structure (yellow) (PDB: 1ATP). f. Local sequence alignment of VPS15 P-loop against PKA. The P-loop motifs are shown in lighter colors. g. The γ-phosphate is aligned 50 degrees away from in line attack on the location of a hypothetical substrate, indicated in blue. h. The terminal phosphates makes extensive hydrogen bonds with residues from the A-loop and catalytic loop of VPS15. An ordered water molecule is proximal to the γ-phosphate.

The unique presence of an Arg at the gatekeeper position reorients the Mg^2+^-coordinating Asp166 of the VPS15 DFG motif towards Arg103. The rotamer switch in Asp166 repositions the catalytic Mg^2+^ such that is coordinates all three GTP phosphates. The corresponding Mg^2+^ in active protein kinases coordinates only the β- and γ -phosphates ^26^. The VPS15 P-loop possesses two changes relative to active kinases that cripple its ability to function as a phosphate stabilizing motif. In conventional kinases, the glycine residues at position 3 and 6 in the P-loop (Fig. 4f) stabilize the β and γ phosphates in a phosphotransferase competent geometry (Fig. 4g). The replacement of Gly by Thr and Phe at positions 35 and 38 renders the VPS15 P-loop incapable of positioning the γ phosphate for in line attack on a phosphoacceptor. The collective effect of these the loss of the two Gly residues and the reorientation of Asp166 is to shift the orientation of the GTP β and γ phosphate by 50 degrees away from their position in active kinases (Fig. 4g), such that they are no longer poised for in line transfer to a phosphoacceptor. This structure rules out that VPS15 is a GTP-dependent protein kinase, although the structure is compatible with GTP hydrolysis activity.

## Discussion

The results described above have clarified key aspects of the heretofore mystery-shrouded process of PI3KC3 activation. The data illuminate the mechanism of action of RAB proteins in the process. The results show how the conformational landscape of the complex relates to functional activation. We described a previously unobserved activated conformation compatible with lipid kinase activity. The geometry of membrane docking, which is prerequisite for activation, has been clarified. We found that the active conformation makes much more extensive and intimate contact with membranes than anticipated. We discovered an unexpected coupling between membrane docking and enzyme activation that depends on sequestration of the VPS15 *N*-myristate in the inactive conformation and its release to dock onto the membrane in the activated conformation. Finally, in the most unexpected discovery, we found that VPS15^PKD^ is a unique adaptation of a pseudokinase that binds GTP instead of ATP. Finally, we found that the GTP binding site is intertwined closely with sequestration of the *N*-myristate and allosteric regulation of the VPS34 catalytic site.

RAB1A^20^ and its yeast ortholog Ypt1^27^ are implicated in autophagy initiation, while RAB5 is fundamental to early endosome maturation^12,28^. The structure reported here reveals the RAB1A:VPS34 interface in high-resolution detail, including ordered water molecules. Interface features are conserved between RAB1A and RAB5, which is consistent with the observation that RAB1A interacts directly only with the VPS34 subunit that is common to both PI3KC3-C1 and - C2 complexes. The differential specificity of RAB1A for PI3KC3-C1 and RAB5 for -C2 must therefore be attributed to context and conformation, rather than to direct subunit-specific interfaces. The extensive interactions of the GTP-dependent switch regions of RAB1A explain the dependence of binding on the GTP-bound state. RAB1A binding leads to a long-range structural contraction of the complex. A single molecule study of PI3KC3-C2 activation concluded that RAB5 both recruited and allosterically activated PI3P production^19^. We interpret the combined single molecule and cryo-EM results as follows. The long-range contraction in the PI3KC3-C1 structure shifts the equilibrium between the various conformational states to favor the activated state. Nevertheless, most of the RAB1A-bound sample as observed here by cryo-EM is in the inactive and transitional conformations. Therefore, at least for PI3KC3-C1, full population of the active conformation requires additional activating stimuli. The revised model for membrane docking indicates that RAB1A is positioned very close to the membrane. The role of RAB1A as a membrane recruiter and partial allosteric activator poises the complex for further activation by additional events such as engagement with the ULK1 complex^21^ and phosphorylation by ULK1^29^.

The new high-resolution data make it possible to functionally interpret past lower resolution structures. It is now clear that previously reported NRBF2-^17^ and liposome-bound^20^ complexes correspond to the inactive conformation. Previous observations of the importance of VPS34^KD^ mobility^14,16^ relative to the inactive conformation are consistent with the need for a 140° rotation of VPS34^KD^, completely breaking all contacts with VPS15^PKD^, and then forming new ones, so as to reach the active conformation. The liposome-bound inactive structure^20^ likely represents an on-pathway state that has been recruited to a membrane by RAB binding and the BECN1 BARA aromatic finger^30^, and is thereby primed for the activating transition described here. Engagement of the BECN1 β sheet-1 peptide^18^ with the membrane would be a late step in the pathway, subsequent to the activating transition and concurrent with the extended membrane docking mode shown in Fig. 3. The MD simulations shed light on how VPS34 accesses its membrane-bound PI substrate. The DFG motif of the VPS34 A-loop and DRH motif of the catalytic loop cooperate to stably coordinate the ATP γ-phosphate in an orientation that is primed for a phosphotransferase reaction. The reaction center is roughly 5 Å above the glycerophosphates of the lipids, almost exactly the distance spanned by a single inositol headgroup. The PI substrate lipid then diffuses into the VPS34 substrate cleft, where it is phosphorylated by the optimally oriented ATP molecule.

VPS15 has been known, since its discovery, to be myristoylated^11,23^, yet the function of this myristoylation has not been experimentally addressed. Here, we found that the VPS15 *N*-myristate is sequestered in the N-lobe of VPS15^PKD^ and stabilizes the inactive conformation of the PI3KC3 complex via a web of interactions with the VPS15 ^PKD^ P and A-loops and the VPS34 catalytic and A-loops. Thus, the long-ignored *N*-myristoyl modification of VPS15 has emerged as fundamental to PI3KC3 regulation. A myristoyl GTP switch controls the membrane binding of ARF GTPases. In the ARF system, the myristate is buried in an internal groove in the GDP-bound state, and ejected upon conversion to the GTP state^31,32^. This promotes membrane localization of GTP-bound ARF proteins. In the case of ARF GTPases, GDP stabilizes *N*-myristate burial and GTP promotes *N*-myristate release. In the structure presented here, the converse applies in that GTP binding to VPS15^PKD^ stabilizes the web of interactions that sequesters the VPS15 *N*-myristate.

Pseudokinases, which are catalytically inactivated members of the protein kinase superfamily, represent about one tenth of the human kinome^33^. A subset of pseudokinases bind nucleotides with varying degrees of specificity^33^, but none are known to bind specifically to GTP. PI3KC3-C1 purified from HEK 293 cells is GTP bound in our inactive cryo EM structure without the resupply of GTP in any purification buffer, indicative of low off and hydrolysis rates, similar to many small GTPases. The ability of VPS15 to bind to GTP is conferred by the unique and conserved presence of an Arg at the gatekeeper position that dictates nucleotide specificity. This Arg not only switches purine base specificity from adenine to guanine, it also triggers a rearrangement of the catalytic site by forcing a rotamer change on the Mg^2+^-binding Asp of the kinase DFG motif. This in turn repositions the γ-phosphate such that in line attack on an external phosphoacceptor becomes impossible. This coupling between the determinant for purine specificity and γ-phosphate positioning explains why evolution has produced no known GTP-specific protein kinases. GTP binding to VPS15^PKD^ clearly stabilizes the *N*-myristate-in and VPS34^KD^ inactive conformations on the basis of the high-resolution structure of the inactive conformation of PI3KC3-C1. VPS34^KD^ appears to be more mobile in the active conformation, given that the interactions with VPS15^PKD^ are broken and the stabilizing *N*-myristate is ejected. VPS15^PKD^ is also more mobile, which is expected given the looser nature of the contacts with VPS34^KD^. The local resolution of VPS15^PKD^ is therefore lower in the active conformation and it has not been possible to identify the bound nucleotide in the active form with certainty. A major frontier going forward will be to characterize this newly discovered GTP nucleotide binding cycle using the tools of the GTPase field.

Therapeutic enhancement of autophagy and lysosome biogenesis is a major yet elusive goal, that is thought to hold promise for currently untreatable neurodegenerative diseases and other aging-associated diseases^34^. Here, we resolved the active conformation of one of the rate-limiting signaling complexes in autophagy initiation (PI3KC3-C1) and a key player in both autophagy and lysosome biogenesis via endosome maturation (PI3KC3-C2). We identified two previously unknown small molecule binding sites, those for GTP and myristate. Both the GTP-bound and myristate-in states of these respective sites are associated with the inactivated conformation. Therefore, antagonizing either GTP or myristate binding, if it can be done without loss of stability, would be an attractive path for therapeutic autophagy enhancement through allosteric activation of PI3KC3.

## Methods

### Plasmid construction

The sequences of all DNA encoding the PI3KC3-C1 subunits and VPS15 mutants used in this study were codon optimized, synthesized, and then subcloned into the pCAG vector using either restriction digestion or Gibson assembly (New England Biolabs enzymes). ATG14, mCherry-ATG14, BECN1, and VPS34 were untagged, while VPS15 and mutants were tagged with a TSF tag at the C-terminus. All constructs were verified by Sanger sequencing. Details are shown in Table 1.

### Protein expression and purification

To produce PI3KC3-C1, 1 L of HEK GNTi cells were grown to a concentration of 2.0-2.2 × 10^6^ cells/ml. The pCAG VPS15-TSF, pCAG VPS34, pCAG BECN1, and pCAG ATG14 or mCherry-ATG14 plasmids were co-transfected at a 1:1:1:1 mass ratio using the polyethelenimine (PEI) (Polysciences) transfection method. After 48 hours, the cells were pelted at 2200 RPM by centrifugation. The pellet was washed with phosphate buffered saline (PBS), spun again at 500x g for 10 min in a tabletop centrifuge, flash frozen and stored at −80 C for later use. Cell pellets were thawed at room temperature and resuspended in lysis buffer containing 25 mM 4-(2-Hydroxyethyl)piperazine-1-ethanesulfonic acid (HEPES) pH 7.5, 200 mM NaCl, 2 mM MgCl_2_, 25 mM tris(2-carboxyethyl)phosphine (TCEP) and 10% Glycerol. An EDTA-Free Protease inhibitor tablet (Thermo Scientific) was added, and the gently resuspended pellet was transferred to a Pyrex Dounce homogenizer. The cells were Dounce homogenized 30 times. The homogenate was transferred to a new tube and Triton X-100 was added to the cells for a final 1% concentration, gently mixed, and then left to rock at 4 C for 1 hour. Following detergent lysis, the cells were pelleted by centrifugation (17,000 × rpm for 45 minutes at 4 C). The supernatant was mixed with 5 mL Strep-Tactin Sepharose (IBA Lifesciences) at 4 C overnight. The following day, the resin was washed with lysis buffer lacking Triton X-100 until the post column flow was free of protein as measured by Bradford assay. The bound protein was then eluted with 1 mg/mL or 4 mM desthiobiotin, spin concentrated, and immediately injected over a Superose 6 increase 10/300 GL column in 25 mM HEPES pH 7.5, 150 mM NaCl, 2 mM MgCl_2_, 25 mM TCEP. Protein was concentrated aliquoted into 25 μL fractions, flash frozen by liquid nitrogen, and stored at −80 C until used for cryo-EM or other assays.

The GTPase deficient RAB1A(Q70L) protein was produced using the BL21 strain of *E. coli*. The pET vector containing 6x-His-RAB1A(Q70L) was transformed into *E. coli* by heat shock, and plated on ampicillin resistant plates. A single colony was grown at 37 C overnight at a culture volume of 10 mL, before expansion into 1 L culture. The 1 L culture was grown at 37 C with 220 rpm shaking until an OD value of ∼0.6. The flask was quenched with an ice bath for 15 minutes. Protein expression was induced with 0.1 mM Isopropyl β-D-1-thiogalactopyranoside (IPTG), and the cells were grown overnight ∼16 hours at 18 C with shaking at 180 rpm. The cells were harvested at 4000 rpm in a BECKMAN coulter centrifuge, washed 1x with 1 M PBS, and pelleted again at 4000 × g before being flash frozen and stored at −80 C until purification. The frozen pellet was thawed to room temperature before resuspending in 25 mM HEPES pH 7.5, 300 mM NaCl, 2 mM MgCl_2_, 1 mM TCEP and adding a EDTA free protease inhibitor tablet (Thermo Fisher) and phenylmethylsulfonyl fluoride (PMSF) to a concentration of 1 mM. The cells were then lysed by sonicated using a 5 s on 5 s off cycle at 50% power for 6 minutes of cumulative on time. The lysate was clarified by centrifugation (17,000 rpm, 45 minutes). The supernatant was immediately applied to a Ni-NTA column 3x, and washed until the column wash was free of protein by Bradford assay. The protein was eluted using wash buffer containing 300 mM imidazole, concentrated, and applied to an S75 10/300 column 25 mM HEPES pH 7.5, 150 mM NaCl, 1 mM MgCl_2_, 1 mM TCEP. Protein was concentrated to 50 μM, separated into 50 μL aliquots and flash frozen in liquid N_2_ for later use.

### VPS34 lipid kinase assay

VPS34 kinase activity was assessed via the ATP Glo (Promega) luminescence assay. Small unilamellar vesicles (SUVs) were prepared by first dehydrating 0.4 mg of 40% dioleoyl phosphatidylcholine (DOPC), 20% dioleoyl phosphatidylethanolamine, 20% dioleoyl phosphatidylserine, 20% liver phosphatidylinositol (all lipids were from Avanti Polar Lipids) in a glass tube with a gentle N_2_ stream. The lipids were dried overnight in a vacuum dessicator to remove excess chloroform. The following day, the lipids were rehydrated in 25 mM HEPES pH 7.5, 200 mM NaCl, and vortexed to resolubilize. The lipids were repeatedly fractured by 10 cycles of freeze thaw using liquid nitrogen and 37° C water bath. The lipids were sonicated using a small probe tip sonicator at 50% power in repeated 2 seconds on/ 2 seconds off cycles for 15 minutes on ice and water mixture. The SUVs were spun in the benchtop centrifuge and used immediately. A kinase dilution buffer consisting of 25 mM HEPES pH 7.5, 200 mM NaCl,10 mM MgCl2, 1 mM MnCl_2_, 2 mM TCEP, and a 2x lipid dilution buffer consisting of 25 mM HEPES pH 7.5, 200 mM NaCl, 20 mM MgCl_2_, 2 mM MnCl_2_, 4 mM TCEP was prepared. 9 μL 50 nM PI3KC3-C1 was added, along with 9 μL of 0.1 mg/mL lipids in the dilution buffer, and 2 μL 500 μM ATP in the same buffer. The reaction was allowed to proceed for 1 hour at room temperature, before 20 μL of the ADP glo ATP depletion reagent was added. The detection reagent was allowed to react for 40 minutes before the luminescent output was measured for each condition tested.

### HPLC analysis of bound nucleotides

Ion-pair reversed-phase acetonitrile gradient HPLC was used to assay the nucleotide bound to VPS15 with a protocol adapted from ref. ^35^. First, PI3KC3-C1 was denatured by heating in a heat block at 90 C for 10 minutes. The precipitated PI3KC3-C1 were spun at 21000 rpm for 15 minutes. The supernatant was transferred to a new tube. 60 μL of 10 μM ATP, ADP, GTP, GDP, and the supernatant from the protein sample were eluted in a linear gradient of Buffer A: 100 mM KH2PO4, 5 mM tetrabutylammonium bromide (TBA-B), pH 6.0, 1% acetonitrile (ACN) and Buffer B: 100 mM KH2PO4, 5 mM TBA-B, pH 6.0, 30% ACN. The total observation time was 30 minutes per run, with wavelength observation set at 254 nm. The retention times for each eluted nucleotide were then compared to assess the identity of the bound nucleotide.

### Mass Spectrometry

PI3KC3-C1(VPS15-TSF) and PI3KC3-C1(mCherry-ATG14|VPS15-TSF) were buffer exchanged into 0.5 M ammonium acetate using a 5 mL PD-10 (Cytiva) column. The buffer-exchanged protein sample was denatured at 90 °C for 10 minutes, followed by a 15 min spin at 21,000xg to separate out precipitated protein. The final concentration of nucleotide was assessed by Nanodrop A260 absorbance to be approximately 5 μΜ for the mCherry-labeled sample and 1 μΜ for the non-fluorescent protein complex. Nucleotide samples were analyzed using a liquid chromatography (LC) system (1200 series, Agilent Technologies, Santa Clara, CA) that was connected in line with an LTQ-Orbitrap-XL mass spectrometer equipped with an electrospray ionization (ESI) source (Thermo Fisher Scientific, Waltham, MA). The LC system contained the following modules: G1322A solvent degasser, G1311A quaternary pump, G1316A thermostatted column compartment, and G1329A autosampler unit (Agilent). The LC column compartment was equipped with an Ultra C18 column (length: 150 mm, inner diameter: 2.1 mm, particle size: 3 µm, catalog number: 9174362, Restek, Bellefonte, PA). Ammonium acetate (≥98%, Sigma-Aldrich, St. Louis, MO), methanol (Optima LC-MS grade, 99.9% minimum, Fisher, Pittsburgh, PA) and water purified to a resistivity of 18.2 MΩ·cm (at 25 °C) using a Milli-Q Gradient ultrapure water purification system (Millipore, Billerica, MA) were used to prepare LC mobile phase solvents. Mobile phase solvent A was water and mobile phase solvent B was methanol, both of which contained 10 mM ammonium acetate. The elution program consisted of isocratic flow at 0.5% (volume/volume) B for 2 min, a linear gradient to 99.5% B over 1 min, isocratic flow at 99.5% B for 4 min, a linear gradient to 0.5% B over 0.5 min, and isocratic flow at 0.5% B for 17.5 min, at a flow rate of 150 µL/min. The column compartment was maintained at 40 °C and the sample injection volume was 20 µL. External mass calibration was performed in the positive ion mode using the Pierce LTQ ESI positive ion calibration solution (catalog number 88322, Thermo Fisher Scientific). Full-scan, high-resolution mass spectra were acquired in the positive ion mode over the range of mass-to-charge ratio (*m*/*z*) = 300 to 1000 using the Orbitrap mass analyzer, in profile format, with a mass resolution setting of 60,000 (at *m*/*z* = 400, measured at full width at half-maximum peak height, FWHM). Data acquisition and analysis were performed using Xcalibur software (version 2.0.7, Thermo Fisher Scientific).

PI3KC3-C1(mCherry-ATG14|VPS15-TSF) was mixed into a final buffer containing 8 M urea, 50 mM tris(hydroxymethyl)aminomethane (TRIS) pH 8.0, 10 mM TCEP. Added iodoacetamide to the mixture for a final concentration of 15 mM iodoacetamide and incubated at room temperature for 20 minutes. The trypsin digestion commenced with the addition of 8 μL Gold Trypsin (0.5 mg/mL Promega V5280) along with 160 μL of 50 mM Tris pH 8 and 2 μL 100 mM CaCl_2_. The final buffer composition was 110 mM ammonium acetate, 50 mM TRIS pH 8, 1 mM CaCl_2_, and 2 mM TCEP. The digestion proceeded at 37 °C overnight with shaking at 200 rpm. The following morning, digestion was confirmed by SDS-PAGE, as the bands corresponding to the PI3KC3-C1 components were not present after digestion. Samples of trypsin-digested proteins were analyzed using the 1200 series LC system and LTQ-Orbitrap-XL mass spectrometer (Agilent and Thermo Fisher Scientific, respectively). The LC column compartment was equipped with a Zorbax 300SB-C8 Micro Bore Rapid Resolution column (length: 150 mm, inner diameter: 1.0 mm, particle size: 3.5 µm, part number: 863630-906, Agilent). Acetonitrile, formic acid (Optima LC-MS grade, 99.9% minimum, Fisher, Pittsburgh, PA), and water purified using the Milli-Q Gradient system (Millipore, Billerica, MA) were used to prepare LC mobile phase solvents. Solvent A was 99.9% water/0.1% formic acid and solvent B was 99.9% acetonitrile/0.1% formic acid (volume/volume). The elution program consisted of isocratic flow at 1% (volume/volume) B for 2 min, a linear gradient to 35% B over 90 min, a linear gradient to 95% B over 1 min, isocratic flow at 95% B for 6 min, a linear gradient to 1% B over 1 min, and isocratic flow at 1% B for 20 min, at a flow rate of 120 µL/min. The column compartment was maintained at 50 °C and the sample injection volume was 10 µL. Full-scan, high-resolution mass spectra were acquired in the positive ion mode over the range of *m*/*z* = 340 to 1800 using the Orbitrap mass analyzer, in profile format, with a mass resolution setting of 60,000 (at *m*/*z* = 400, FWHM). In the data-dependent mode, the ten most intense ions exceeding an intensity threshold of 10,000 raw ion counts were selected from each full-scan mass spectrum for tandem mass spectrometry (MS/MS) analysis using collision-induced dissociation (CID). MS/MS spectra were acquired using the linear ion trap, in centroid format, with the following parameters: isolation width 3 *m*/*z* units, normalized collision energy 28%, default charge state 3, activation Q 0.25, and activation time 30 milliseconds. Real-time charge state screening was enabled to exclude singly charged ions and unassigned charge states from MS/MS analysis. To avoid the occurrence of redundant MS/MS measurements, real-time dynamic exclusion was enabled to preclude re-selection of previously analyzed precursor ions, with the following parameters: repeat count 2, repeat duration 10 s, exclusion list size 500, exclusion duration 30 s, and exclusion mass width ±10 parts per million. Raw data files were searched against the amino acid sequences of mCherry-ATG14 Class III phosphatidylinositol 3-kinase complex I (mCherry-PI3KC3-C1) proteins using Proteome Discoverer software (version 1.3, SEQUEST algorithm, Thermo Fisher Scientific) for tryptic peptides (i.e., peptides resulting from cleavage C-terminal to arginine and lysine residues, not N-terminal to proline residues) with up to three missed cleavages, with carbamidomethylcysteine as a static post-translational modification and N-terminal acetylation, N-terminal myristoylation, phosphoserine and phosphothreonine as dynamic post-translational modifications. Assignments were validated by manual inspection of MS/MS spectra.

### Sample preparation of PI3KC3-C1∼RAB1A for CryoEM

RAB1A(Q70L) was loaded with GTP by first stripping protein of its bound divalent cations with 5 mM EDTA at room temperature for 30 minutes. A large molar excess 50 mM GTP was added to the protein and incubated for 30 minutes. The reaction was then quenched with 20 mM MgCl_2_ at room temperature for 20 minutes. The resulting GTP loaded RAB1A was desalted with a PD-10 column in a final buffer of 25 mM HEPES pH 7.5, 150 mM NaCl, 2 mM MgCl_2_, 25 mM TCEP. The PI3KC3-C1 protein was then mixed with RAB1A at a 1:20 molar ratio (0.55 μM PI3K:11 μM RAB1A) and incubated for 1 hour at room temperature. This sample was used immediately for grid preparation.

### Sample vitrification and data acquisition

For cryo-EM grid preparation, n-Octyl-β-D-Glucopyranoside (OG) was added to the 1:20 molar ratio PI3KC3-C1:RAB1A at a final concentration of 0.05%. 3 μL of the resulting sample was immediately applied onto freshly glow-discharged (PELCO easiGlow 25 mA current, 1 min) QUANTIFOIL R 2/1 mesh Cu 300 holey carbon grids. Sample vitrification was accomplished with a Vitrobot cryo-plunger (Thermo Fisher Scientific) using 100% humidity, 4 C, 3 s wait time, −15 blot force. The dataset of the PI3KC3-C1-RAB1A complex were recorded on a 300 kV Titan Krios microscope equipped with an X-FEG and energy filter set to 20 eV. The data was automatically collected with SerialEM on a K3 Summit direct electron detector (Gatan) at a 81,000x magnification in super resolution pixel size of 0.525Å and a defocus range of −0.8 to - 2.2 micrometers. 50 frame image stacks were collected to a final cumulative dose of ∼50 e/ Å^2^.

### Image processing and 3D reconstruction

The data sets were processed in cryoSPARC v3^36^. The super resolution movies were motion corrected and fourier cropped 2x using the cryoSPARC implementation of Patch Motion Correction^37^. Contrast transfer function determination was done using Patch CTF Estimation in cryoSPARC v3. Single particles from 10 micrographs across a range of defocus values were manually picked and used to train a Topaz ^38^ model. The particles were extracted with a box size of 400×400×400 or ∼1.5x the diameter of a single PI3K complex to ensure retention of delocalized CTF information, and binned 4x to increase computational speed for particle sorting. Next, two-dimensional classification was applied to the extracted particles, and obvious junk particles were excluded from subsequent processing. Next, a junk class from the early rounds of an ab initio run, plus a map generated using an apo PI3K model in UCSF Chimera using molmap at a resolution of 20 A were used in heterogeneous refinement for 3-4 rounds until a healthy substack of particles was evident by 2D classification. Finally, a three class *ab initio* reconstruction was used for a final particle clean up, and produced a healthy model that showed obvious density for RAB1A. The particles in the healthy class were re-extracted at a full 400 pixel box size, and homogeneous refinement was performed using the *ab-initio* model and the clean particle stack to generate a high resolution model and particle alignments for downstream classification. To further classify the particles, 3D classification without alignment using a large mask on the putative kinase domain and 50 classes was performed in cryoSPARC. The outputs from this job generated several strongly populated classes that showed distinct conformations for the kinase domain. The three obvious classes from the group of 50 were then used as input for a heterogeneous refinement job. Each class was populated with non-uniform probability, indicating that the classification was successful. The first and largest class of particles was subjected to non-uniform refinement. Local refinement was performed on three parts of the complex to improve features on the particle periphery, including the VPS15 pseudokinase domain, the RAB1A interface region, and the BECN1/ATG14 BARA dimer domain. Masks were created in UCSF ChimeraX v1.5 using Volume Tools. Each population that showed strong density for the kinase domain was subjected to 3D-variability analysis in cryoSPARC in cluster mode, and the cluster of particles that corresponded to the strongest density for the VPS34 kinase was used in conjunction with the map output from 3D variability analysis for a final round of non-uniform refinement. Local refinement was performed on the kinase domain and each region listed above separately to yield the highest resolution features for each part of the map. We ran local resolution estimation in cryoSPARC on each locally refined region of the intermediate complex using the local refinement masks. The locally refined maps were, for each distinct kinase conformation, aligned in UCSF ChimeraX ^39^ and combined using the vop maximum command, and used for model building and visualization. The details of data processing are summarized in Table 2.

### Cryo-EM model building

A starting model of the full-length PI3KC3-C1 in both the active and inactive state, and RAB1A were generated using the ColabFold implementation of AlphaFold2^40,41^. The resolution of the experimental observations permitted amino acid assignment for the inactive and intermediate maps, while for the active state, the models could be built only at the level of secondary structural features. Iterative manual model building using the MD-based model building software ISOLDE^42^ as implemented in UCSF ChimeraX^39^ was used in conjunction with automated Phenix real space refinement to arrive at an atomic model that closely matched the EM density maps in each case.

The 2.73 Å resolution transitional structure showed both side chain features as well as higher-resolution features such as GTP, myristate density, and water molecules to be placed near the active sites of the VPS15 and RAB1A proteins (Extended data Fig. 4). The water molecules in the RAB1A fold and at the RAB1A-VPS34 interface were compared to existing crystal structure water molecules, and half map occupancy and were modeled in the event of coincident waters in the crystal structures and the map. In the case of VPS15, waters were modeled in using a combination of half map occupancy, Q-score, and whether similar thresholds showed water molecules in the RAB1A region of the reconstruction. The 3.1 Å inactive state map permitted the identifications of side chain features at the interface between VPS15 and VPS34. We could still unambiguously observe the GTP molecule in the inactive map, although the lower resolution did not unambiguously resolve the same ordered water molecules as in the transitional state. We could see direct contact between the VPS34 A-loop and the N-terminal region of VPS15. The active kinase structure was locally refined using a mask that encompassed the VPS34 kinase domain. The VPS34^KD^ showed broad secondary structure features, but we could not unambiguously determine side-chain conformations. As a result, the kinase domain was modeled using a polyalanine, and an AlphaFold2 model of the VPS15-VPS34 subcomplex was used to assess the proximity of key residues at the active interface.

### MD simulations

MD simulations were performed with GROMACS 2020^43^ using the CHARMM36m force field^44^. All-atom structural models of PI3KC3-C1 consisting of full-length VPS34 and VPS15 in complex with BECN1 (with its 102 unstructured N-terminal residues excluded from the model) and ATG14 (excluding unstructured residues 1-38) were prepared based on the active conformation as determined by cryo-EM, with missing regions predicted by AlphaFold-Multimer 2.2^40^. Protonation states of amino acid side chains were assigned according to pK_a_ prediction by PROPKA^45^. A molecule of ATP was parameterized using CHARMM-GUI^46^ and placed into the VPS34 ATP-binding site alongside a coordinating Mg^2+^ ion. Six independent replicates of the simulation system were prepared, beginning with construction of membranes consisting of randomly distributed coarse-grained lipids using the *insane* method^47^, with 60% DOPC, 20% DOPE, 5% DOPS, and 15% POPI. Each membrane patch was solvated with 150 mM of aqueous NaCl, equilibrated for 200 ns, and converted into an atomistic representation using the CG2AT2^48^ tool. Atomistic PI3KC3-C1 was placed above the resulting membranes with a minimum distance of ∼2 nm between protein and lipid atoms. The protein-membrane systems thus produced (with dimensions ∼29 × 29 × 30 nm^3^) was subsequently re-solvated and subjected to 10 ns of further equilibration followed by 2 µs production runs. Harmonic positional restraints, with a force constant of 1000 kJ mol^−1^ nm^−2^, were applied to non-hydrogen protein atoms during equilibration. System pressure and temperature were maintained at 1 bar and 310 K, respectively, using a semi-isotropic Parrinello-Rahman barostat^49^ and the velocity-rescaling thermostat^50^ in the production phase. Long-range electrostatic interactions were treated using the smooth particle mesh Ewald method^51^, with charge interpolation through fourth-order B-splines. The integration time step was 2 fs, and the LINCS algorithm^52^ was used to constrain covalent bonds involving hydrogen atoms. For steered MD simulations, harmonic restraints with a force constant of 100 kJ mol^−1^ were applied to reduce the center-of-mass *z*-distance between the protein group in question (either the Kα12 helix of VPS34 or residues Phe270 and Phe274 of BECN1) and the membrane lipids underneath, at a rate of –0.5 nm ns^−1^, until the protein group was at the membrane surface. The complex was subsequently allowed to relax over periods of 2 µs during further simulations upon removal of any steering force.

## Acknowledgments

We thank members of the Hurley Lab, D. Fracchiolla, and others in Aligning Science Across Parkinson’s (ASAP) Team mito911 for advice and discussions. We thank D. Toso, P. Tobias and R. Thakkar for cryo-EM facility support.

## Data availability

The cryo-EM maps are being deposited in the Electron Microscopy Data Bank (EMDB). Protocols are being deposited in protocols.io. Plasmids developed for this study are being deposited at Addgene.org. Original gel scans, mass spectrometry RAW data files, HPLC data, and ADP glo data are being deposited at Zenodo.

## Author contributions

Conceptualization: J.H.H. Investigation: A.C., M.C., X.R., S.G., S.R., A.C.C., A.I. Visualization: A.C. Supervision: J.H.H., G.H. Writing—original draft: A.C. and J.H.H. Writing—review and editing: All authors.

## Extended Data

**Extended Data Figure 1.**
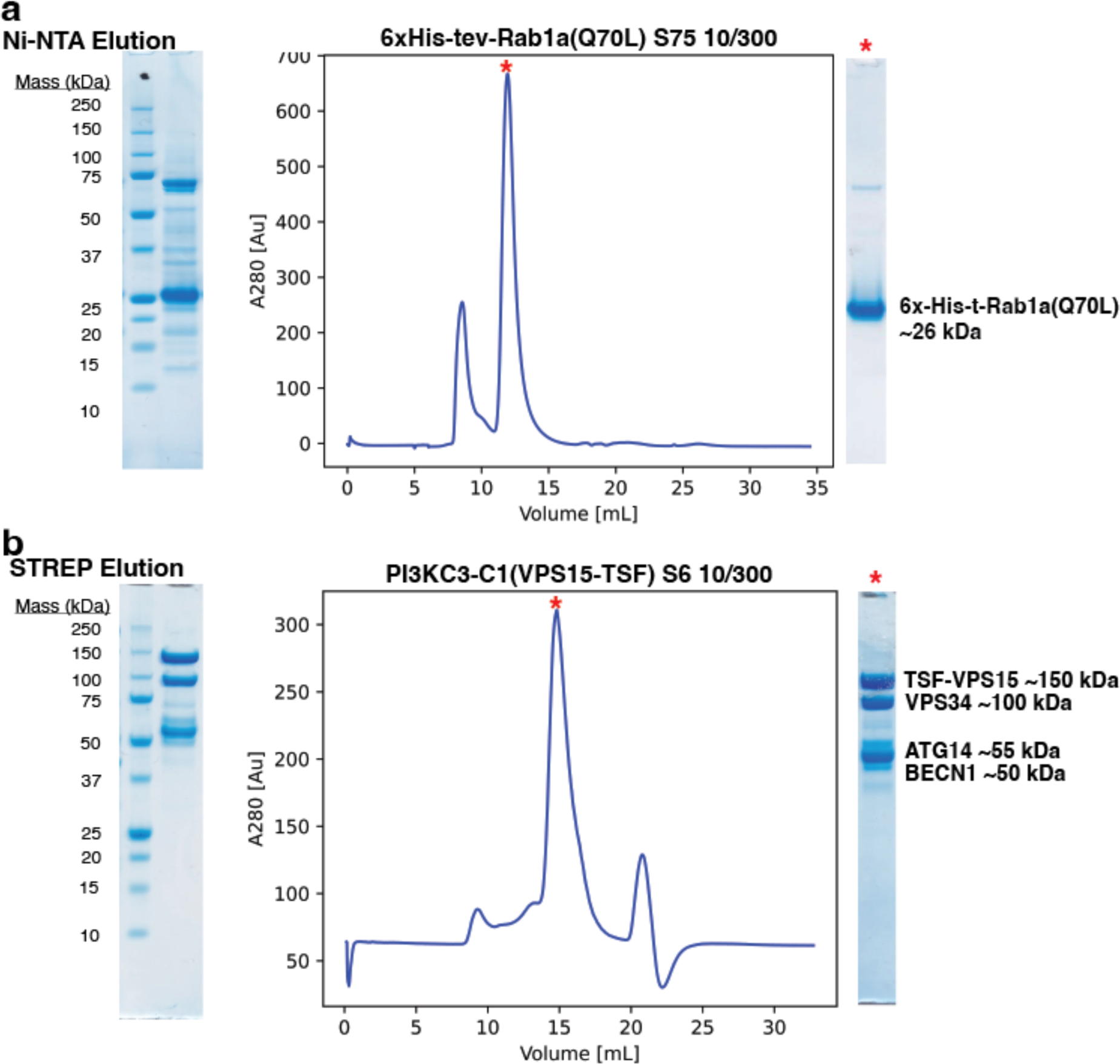
A) Purification of 6xHIS-RAB1A. Shown are the results of Ni-NTA affinity purification, and the result of size exclusion using S75 10/300. Molecular weight is indicated beside the main protein band. B) Purification of PI3KC3-C1 (VPS15-TSF). Show are the results of STREP resin affinity purification size exclusion. The molecular weights of each complex component are shown beside the results from size exclusion.

**Extended Data Figure 2.**
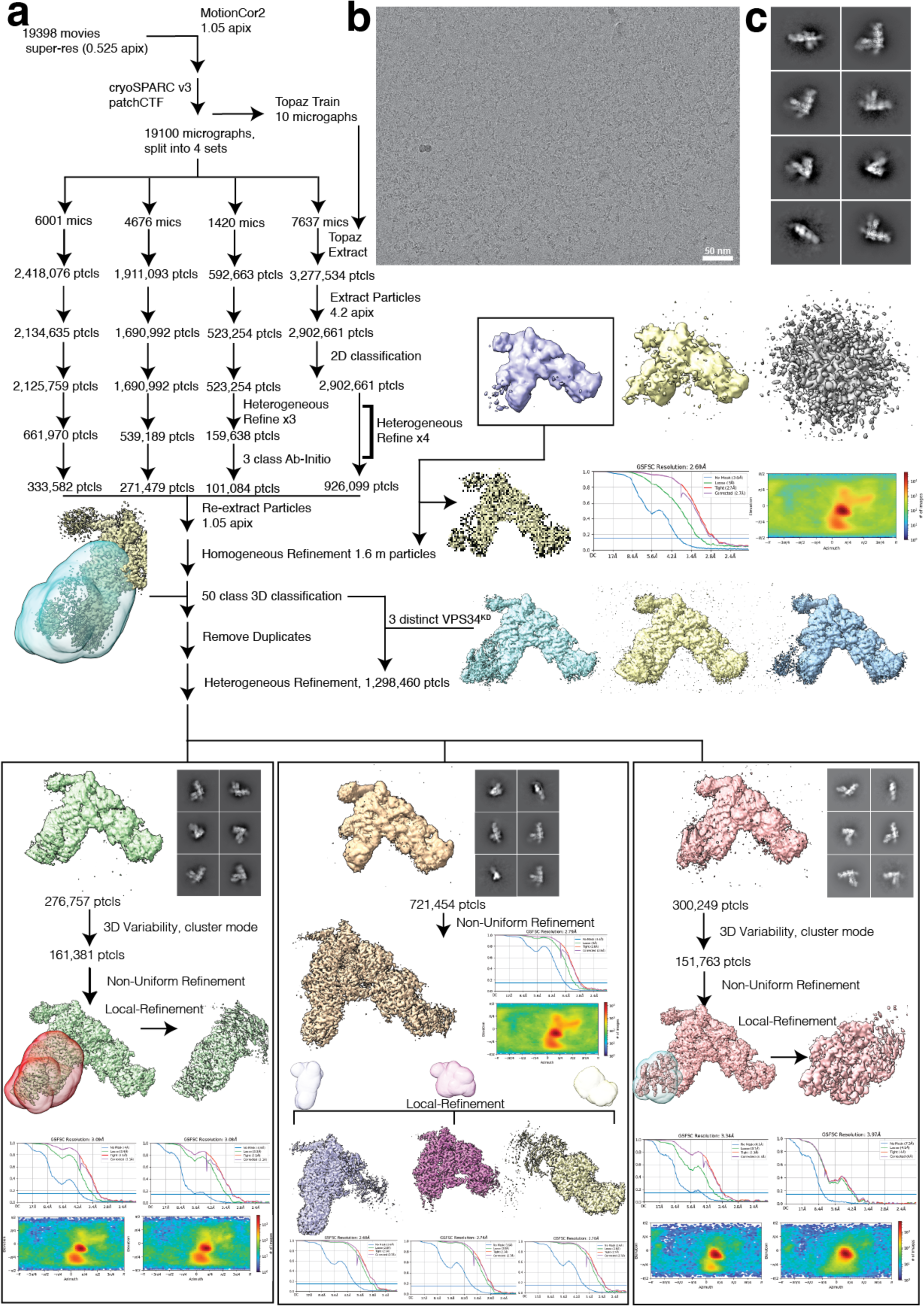
A) Data processing pipeline. Details for the pipeline are in the methods. Briefly, a ∼19,000 movie dataset of PI3KC3-C1 bound to RAB1A was collected and processed in cryosparc using multiple rounds of 2D and heterogeneous refinement to yield a clean particle subset. Further 3D classification using a mask around the VPS34 kinase domain found three distinct classes. Further processing showed two rigid conformations for the kinase domain, and one transitional state. B) Representative motion corrected micrograph. The scale bar indicates 50 nm. C) representative 2D class averages from the initial round of bulk particle sorting. Clear secondary structural features are apparent.

**Extended Data Figure 3.**
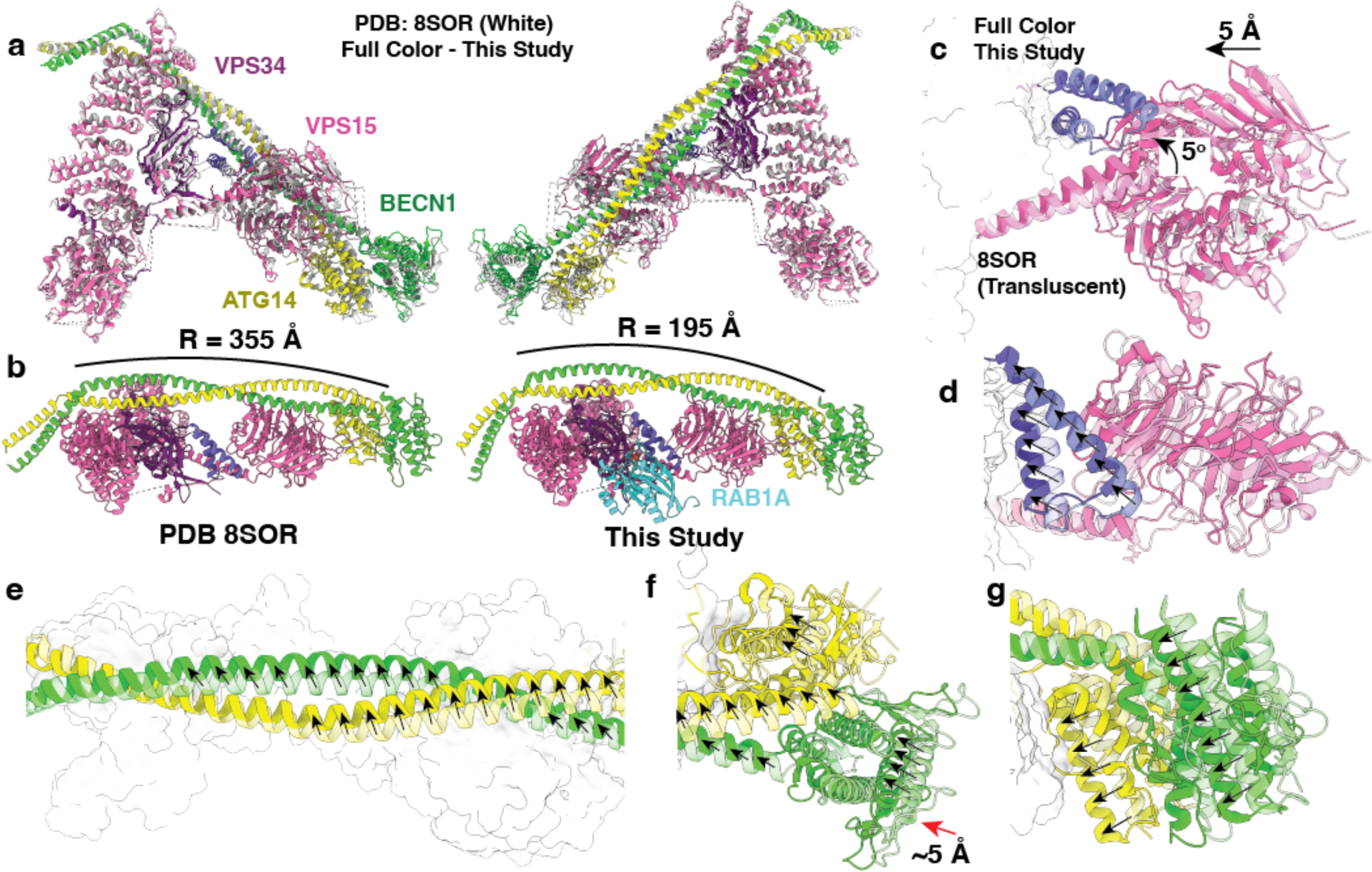
a) A structural alignment of the PI3KC3-C1∼RAB1A complex (colors) to the the Apo PI3KC3-C1 complex (white, PDB: 8SOR). b) The complex changes radius of curvature upon binding RAB1A, going from R = 355 to R = 195 angstroms. c) Conformational change showing the angular displacement of the VPS15 helical linker, and the global contraction of the WD40 domain. d) The VPS34 C2-HH motif shifts inward and upward in response to binding RAB1A. e) The ATG14/BECN1 coiled coil shifts up and rotates about the main axis of the complex, as indicated by the black displacement vectors. f) the contraction fo the BECN1 BARA and ATG14 BARA like domains Is indicated using the vectors. g) Similar to f, from the side view.

**Extended Data Figure 4.**
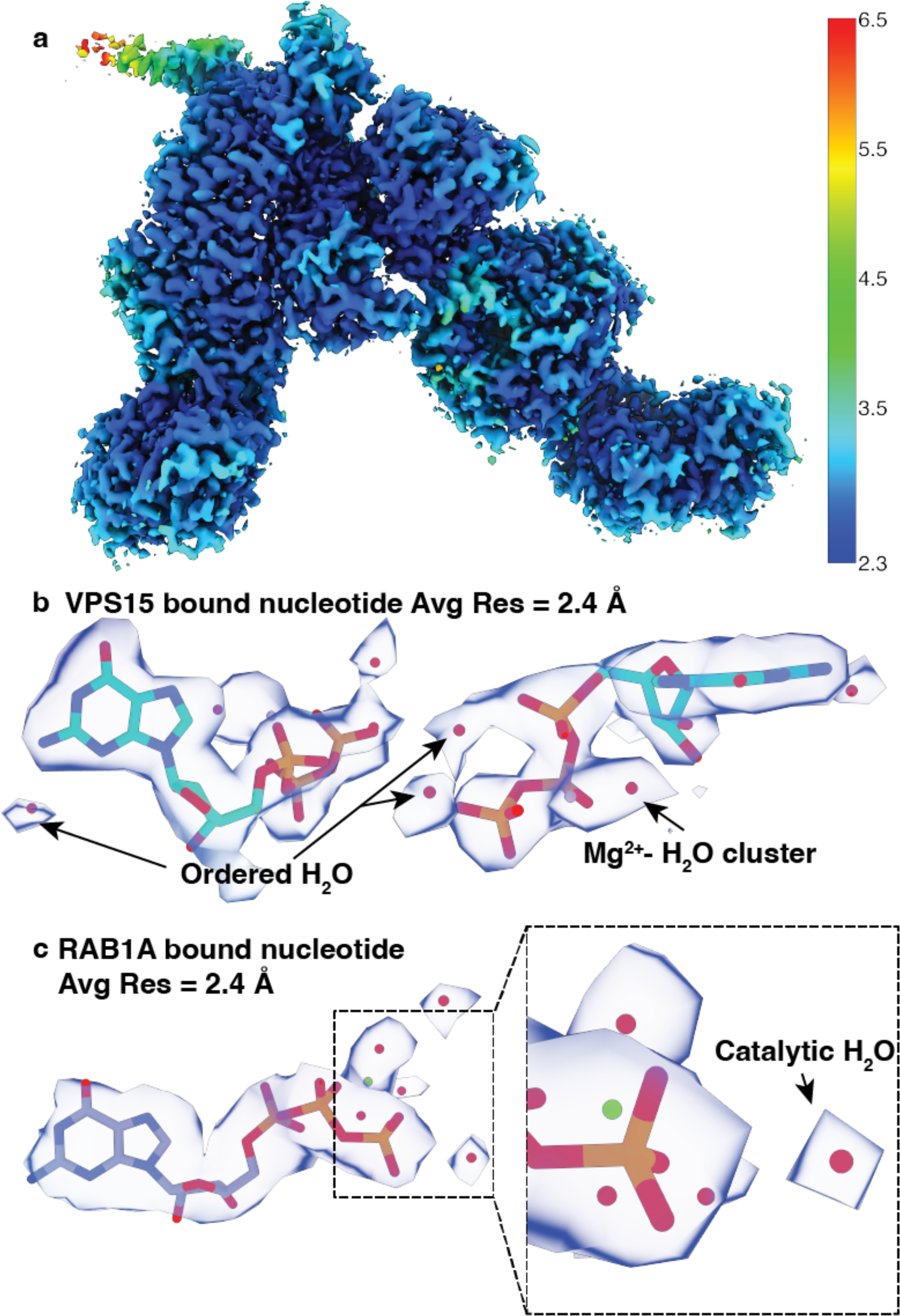
a) Local resolution estimate of the intermediate 2.73 Å structure. The resolution ranges from 2.3 to 2.6 Å in the lowest B-factor regions of the complex, including the VPS15 nucleotide pocket and RAB1A nucleotide binding pocket. b) detailed look a the GTP molecule bound to VPS15. The density clearly shows density non-proteinaceous and non-nucleotide features including a Mg^2+^ cluster, and multiple H_2_O molecules. C) detailed look at the RAB1A reaction center. The Mg^2+^ - H_2_O cluster is apparent in this GTP molecule, as is the hydrolytic water molecule, positioned just downstream of the γ-phosphate.

**Extended Data Figure 5.**
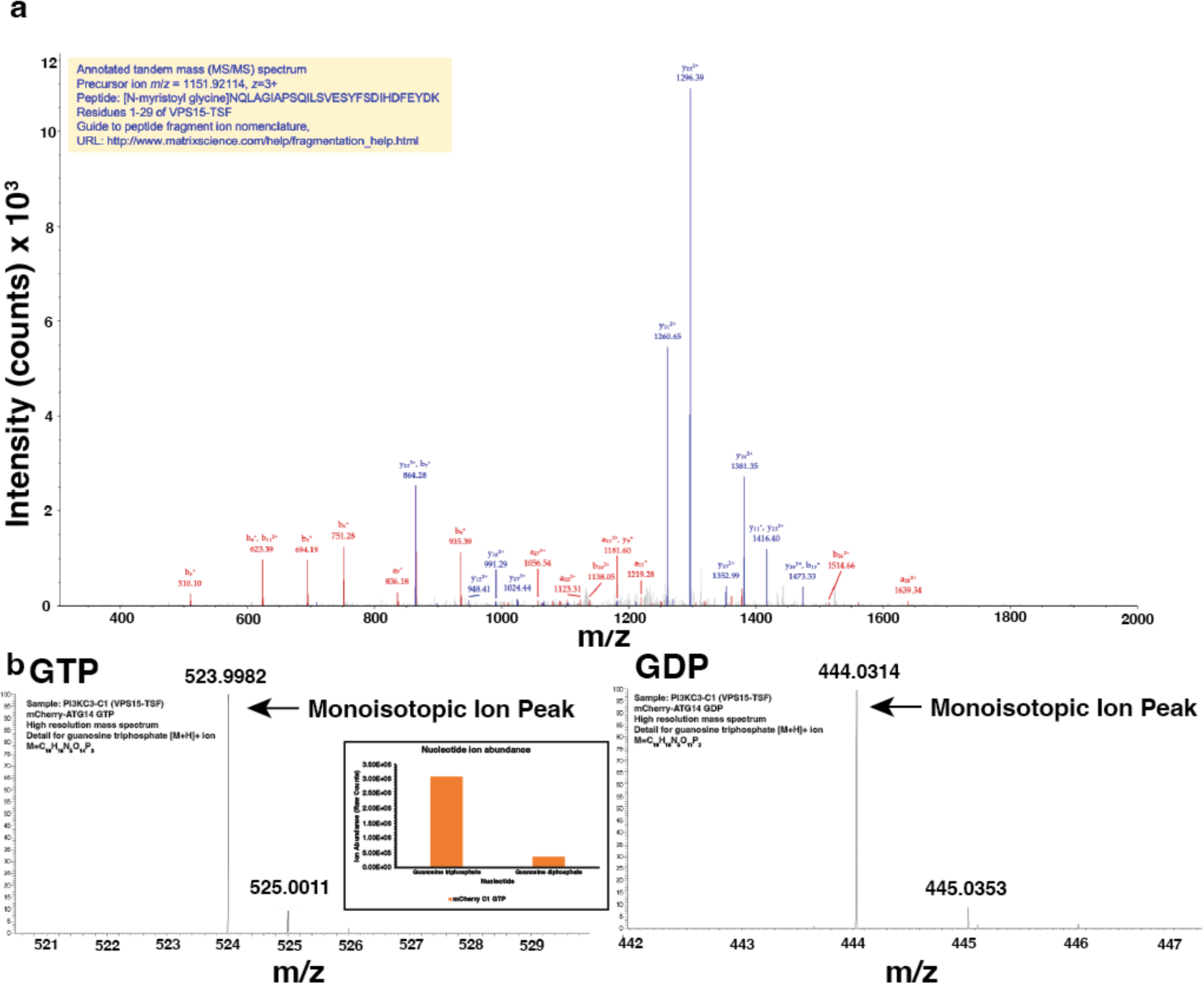
a) Trypsin-digestion coupled to LC-MS/MS was used to resolve the N-myristoyl modification of VPS15. Shown is a tandem mass (MS/MS) spectrum corresponding to the N-terminal 1-29 residues of VPS15. b) Electrospray ionization high-resolution mass spectra of nucleotides that were eluted from a sample of PI3KC3-C1 (VPS15-TSF and mCherry-ATG14). The monoisotopic ion peaks clearly correspond to GTP and GDP in a roughly 10:1 abundance ratio.

**Extended Data Figure 6.**
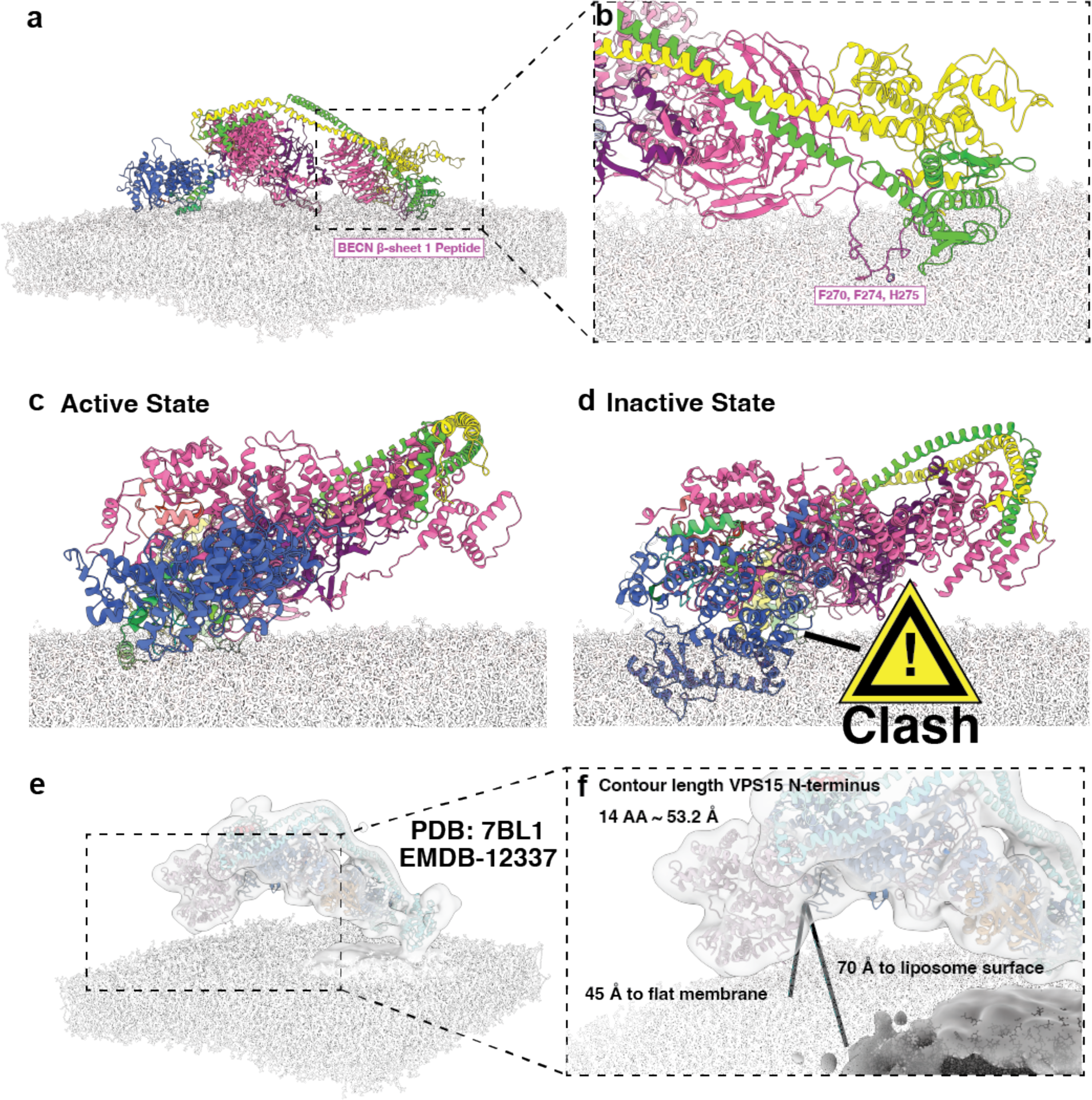
A) Molecular dynamics snapshot of the membrane bound PI3KC3-C1. B) Close up view of the BECN β-sheet 1 unfolded and interacting with the membrane. Putative membrane interactors Phe270, Phe 274, and His275 are indicated in the figure. C) End-on view fo the active complex, showing the intimate membrane contact. D) The same view of the inactive complex showing incompatibility of this orientation with membrane access. E) Stereo view of the RAB5 bound PI3KC3-C2 model (PDB:7BL1) docked into the electron tomography reconstruction, shown without liposome masked out (EMDB:12337). F) A close-up of the VPS34^KD^, indicating the distance to a putative flat membrane (45 Å), and the liposome surface (70 Å). The contour length of 53.2 Å of the VPS15 N-terminal disordered 14 residues is indicated, assuming an average persistence length of ∼3.8 Å per amino acid.

**Extended Data Figure 7.**
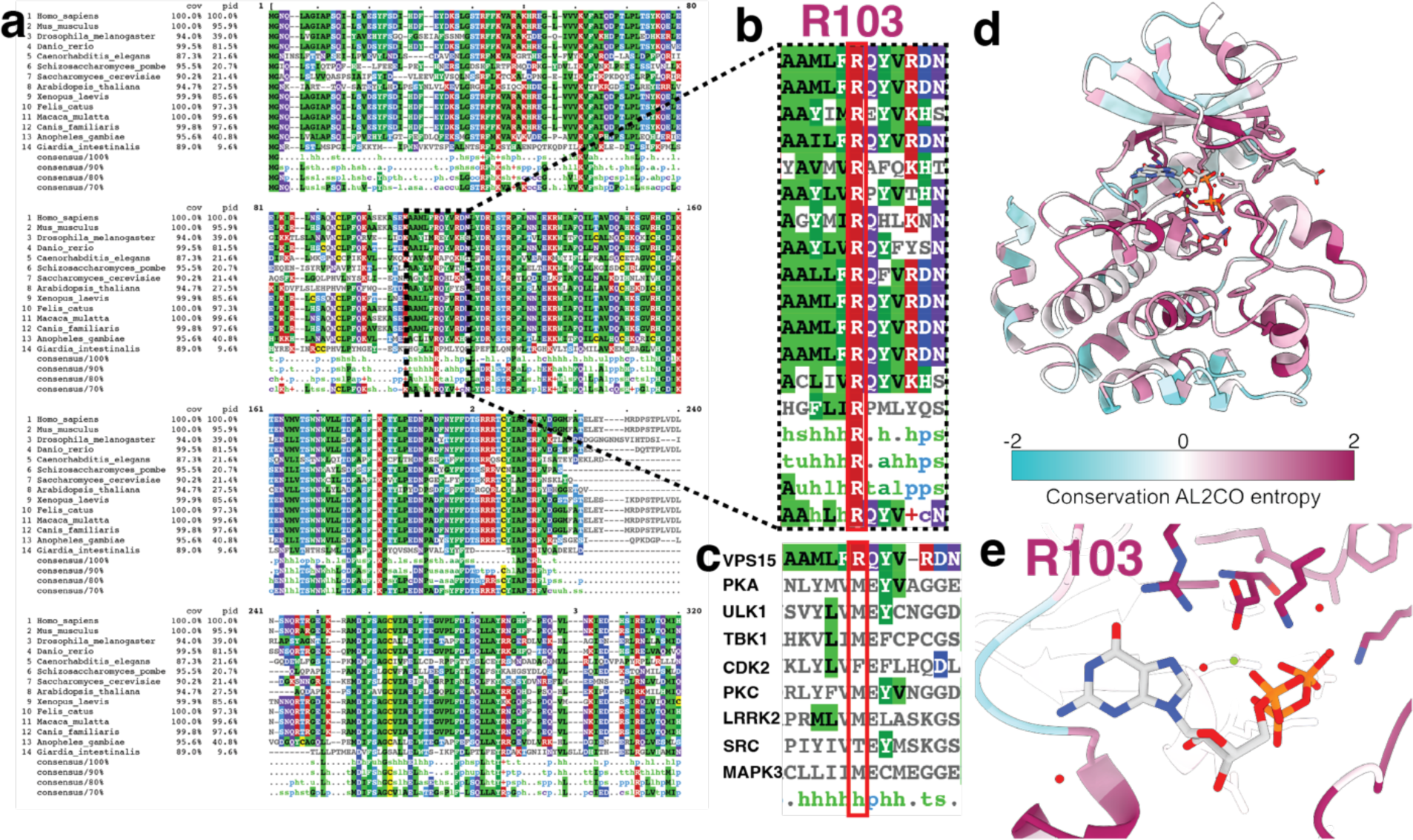
A) A multiple sequence alignment of *<u>Homo sapiens</u>* VPS15 and homologs for multiple model organisms, including *<u>Mus musculus, Drosophila melanogaster,</u> <u>Danio rerio, Caenorhabitis elegans, Scizosaccharomyces pombe, Saccharomyces cerevisiae,</u> <u>Arabidopsis thaliana, Xenopus laevis, Macaca mulatta,</u>* and several other common organisms. Multiple sequence alignments were generated by manual accession from the Uniprot Uniref100 database, and submission to the B) Inset of the gatekeeper residue, showing the absolute conservation of the gatekeeper Arg. C) Multiple sequence alignment of VPS15 with several eukaryotic protein kinases. The consensus hydrophobic amino acid is broadly shared, while VPS15 contains an arginine at this position. D) Sequence conservation coloration of VPS15 based on the AL2CO entropy measure. A multiple sequence alignment of 100 homologs produced using a BLAST search in UCSF ChimeraX was used to calculate the entropy measure. E) A close up of the gatekeeper Arg hydrogen bond interactions with GTP colored by conservation.

